# Dissociation kinetics and avidity gate SARS-CoV-2 neutralization by HR2 stem helix antibodies

**DOI:** 10.64898/2026.08.17.745160

**Authors:** Virginia Crivelli, Concetta Guerra, Morgan E. Abernathy, Jacopo Sgrignani, Giada Zoppi, Mia L. Greeson, Annalisa Sanga, Patrizia Locatelli, Jasmine Cantergiani, Benedetta Cena, Tomás Cervantes Rincón, Yu E. Lee, Michael Eso, David Jarrossay, Maira Biggiogero, Veronica Calvaruso, Alessandra Franzetti Pellanda, Christian Garzoni, Elia Tamagnini, Sara Lestani, Luca Varani, Sina Sommer, Daniel Fernandez, Giovanna Barba-Spaeth, Emily G. Niejadlik, Stylianos Bournazos, Davide F. Robbiani, Christopher O. Barnes, Andrea Cavalli

**Author notes:** These authors contributed equally to this work (Virginia Crivelli, Concetta Guerra, Morgan E. Abernathy, Christopher O. Barnes). **Author contributions:** V.C., C.G., M.E.A., D.F.R., C.O.B., and A.C. designed research; V.C., C.G., M.E.A., J.S., G.Z., M.L.G., A.S., P.L., J.C., B.C., T.C.R., Y.E.L., M.E., D.J., E.T., S.L., L.V., S.S., D.F., G.B.S., E.G.N. and S.B. performed research; A.F.P., M.B., V.Cal., and C.Gar. contributed COVID-19 patient cohort; V.C., C.G., M.E.A., J.S., M.L.G., and D.F. analyzed data; and V.C., C.G., M.E.A., D.F.R., C.O.B., and A.C. wrote the paper with input from all authors. **Competing Interest Statement:** The Institute for Research in Biomedicine has filed a patent application in connection with antibody hr2.016.

## Abstract

SARS-CoV-2 evolution has reduced the efficacy of clinical monoclonal antibodies, underscoring the need for therapeutics targeting conserved viral regions. The Spike (S) heptad repeat 2 (HR2) stem helix is highly conserved across SARS-CoV-2 variants and related betacoronaviruses. Although neutralizing antibodies to this region have been identified, the evolution of humoral responses to HR2 and the determinants of effective neutralization at this site remain poorly understood. We previously identified human neutralizing antibodies to a conserved peptide within this region (HR2 coldspot). Here, longitudinal analysis over 30 months shows that HR2-specific antibodies persist and undergo somatic hypermutation, yet antibodies isolated at later time points did not surpass the breadth or potency of hr2.016, which emerged shortly after primary infection. Crystal structures of four HR2 stem-helix antibodies revealed convergent recognition across distinct antibody lineages. However, comparison of hr2.016 with its non-neutralizing clonal relative hr2.086 showed that adopting this shared binding mode is not sufficient for effective neutralization. Characterization of this antibody pair through surface plasmon resonance and molecular dynamics simulations revealed that robust neutralization requires slow intrinsic dissociation reinforced by avidity. Together, these findings show that continued evolution of HR2-specific responses does not necessarily enhance antibody breadth or potency and that, despite convergent epitope recognition, effective neutralization requires slow intrinsic dissociation reinforced by avidity, highlighting kinetic stability as a key criterion for antibody discovery and vaccine design against viral epitopes.

**Significance statement:** SARS-CoV-2 spike HR2 stem helix is highly conserved and targeted by broadly neutralizing antibodies, but how HR2 responses evolve and what converts its recognition into neutralization remain unclear. We show that HR2 antibodies remain rare after repeated antigen exposure and can accumulate somatic mutations without gaining breadth or potency. Although distinct lineages converge on the same epitope and binding mode, structural similarity does not ensure neutralization. Instead, antiviral activity requires prolonged HR2 engagement, generated by slow intrinsic dissociation and amplified by IgG avidity. These findings establish antibody residence time as a critical determinant of neutralization at the conserved HR2 fusion epitope, providing a kinetic principle to guide antibody discovery and vaccine design.

## Introduction

The clinical efficacy of monoclonal antibodies (mAbs) targeting the SARS-CoV-2 Spike (S) has been repeatedly compromised by viral evolution (1–4). Successive variants of concern (VOCs) rapidly accumulated mutations in the receptor-binding domain (RBD) and N-terminal domain (NTD), leading to widespread escape from authorized mAbs and underscoring the need for antibody therapeutics that target structurally and functionally constrained regions of the trimeric S glycoprotein (5–8).

The heptad repeat 2 (HR2) stem helix within the S2 subunit represents one such region, playing a central role in the large-scale conformational rearrangements required for membrane fusion (8–10). Several human-derived mAbs targeting this region have demonstrated broad neutralizing activity and protection in preclinical models (8, 11–13), highlighting this region as an attractive site of vulnerability. Mechanistically, rather than blocking ACE2 engagement, HR2 stem-helix mAbs interfere with the S2 conformational rearrangements that drive membrane fusion, preventing progression to the post-fusion state (14, 15). Structural studies have further shown that these antibodies, despite arising from distinct donors and recurrent germline lineages, converge on a conserved hydrophobic surface within the stem helix, suggesting that recognition of this epitope is subject to strong structural constraints (16–18). However, the structural determinants that distinguish potent from weakly or non-neutralizing antibodies to this region and their resilience to viral variation remain poorly characterized (8, 12, 16, 19).

We previously reported on a panel of human antibodies derived from unvaccinated individuals after primary SARS-CoV-2 infection (2020–2021) (13) that bind to a short peptide within the HR2 stem helix corresponding to a region with high sequence conservation but low serological prevalence (HR2 coldspot). Among them, hr2.016 emerged as the most potent and broadly reactive neutralizing antibody. However, whether repeated antigen exposure through subsequent infection or vaccination could further increase the breadth or potency of HR2 coldspot-specific responses remained unknown.

By taking advantage of longitudinal cohort samples, we report herein on the serologic evolution of HR2 coldspot IgGs over 30 months, as well as on the clonal persistence and molecular evolution of HR2 coldspot monoclonal antibodies in two individuals. Crystal structures of four HR2 stem helix antibodies revealed convergent recognition of the conserved epitope across multiple germline lineages. Comparison of hr2.016 with its non-neutralizing clonal relative hr2.086 shows that dissociation kinetics and avidity are key determinants of neutralization.

These findings point to a kinetic threshold for neutralization at the HR2 stem helix mediated by bivalent binding, providing a framework for understanding the mechanism of antibody neutralization at this conserved epitope.

## Results

### Evolution of HR2 coldspot antibodies

We previously showed that 6 months after primary infection with SARS-CoV-2, a fraction of convalescent individuals display plasma IgG antibodies to a peptide (amino acid positions 1144-1163 of S) corresponding to a highly conserved region of S located at the HR2 stem helix (HR2 coldspot) (13). To evaluate whether HR2 coldspot antibodies persist and evolve over time, we compared their presence at 6 months with 18 and 30 months in the same individuals (n = 71) (13, 20). As determined by the enzyme-linked immunosorbent assay (ELISA), plasma IgGs to the HR2 coldspot were significantly increased in the cohort at 18 months and subsequently declined at 30 months (Fig. 1A and Fig. S1A).

**Figure 1.**
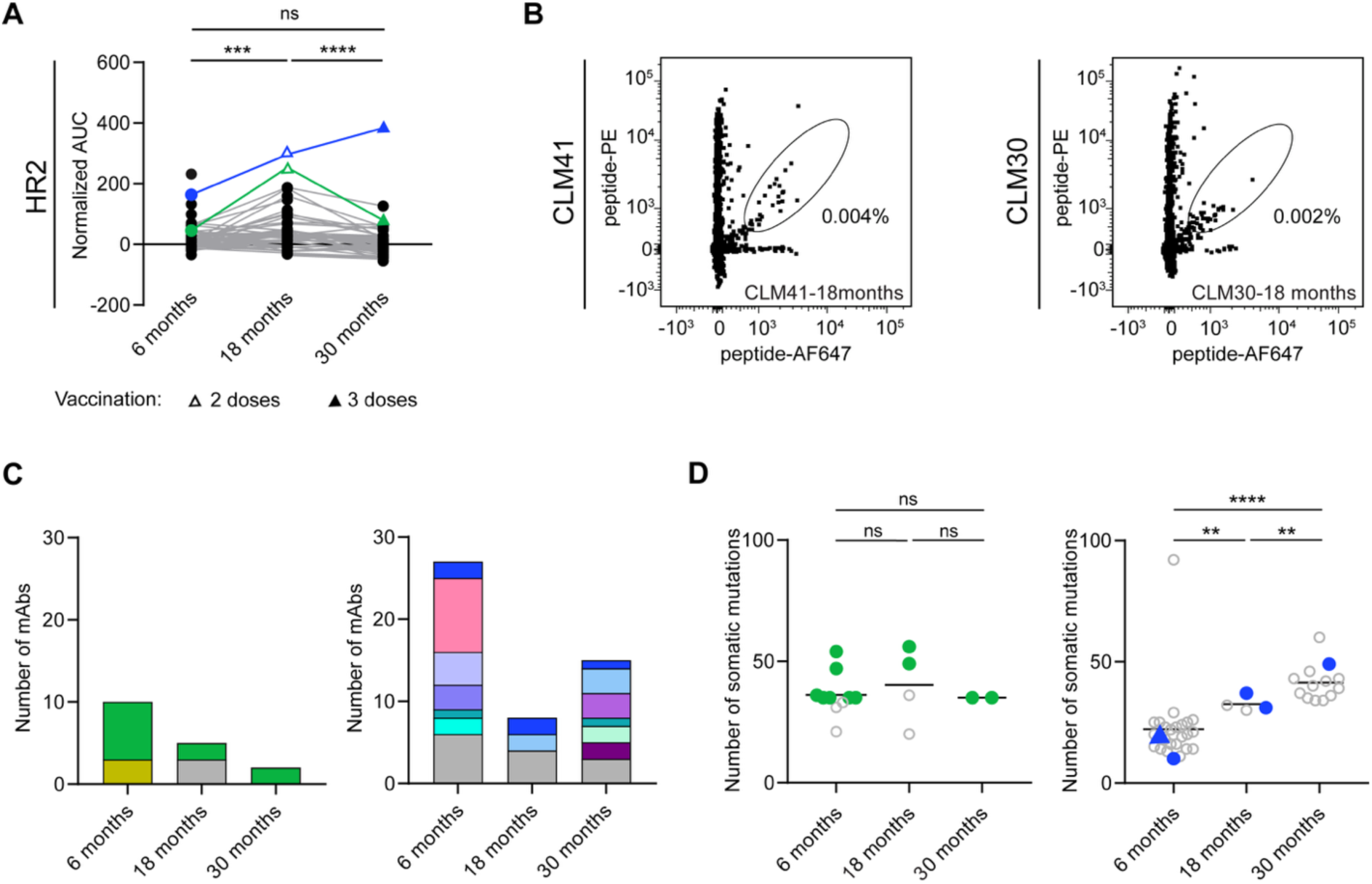
Evolution of HR2 coldspot antibodies. **(A)** Plasma IgG reactivity over time to the HR2 coldspot peptide, represented as normalized area under the curve (AUC) of ELISA measurements. Mean of two independent experiments. Green (CLM41) and blue (CLM30) represent samples from individuals selected for HR2 coldspot antibody analysis. Empty triangle indicates that the individual received two doses of vaccine, filled triangle is at least three doses. Paired Wilcoxon test. *** is p < 0.001 and **** is p < 0.0001 **(B)** Representative flow cytometry plots of B cells binding to HR2 coldspot peptide. The percentage of cells in the sorting gate is shown. **(C)** Clonally related HR2 coldspot antibody sequences over time. Each color represents sequences of related antibodies, except grey, which represents singlets. **(D)** Number of heavy and light chain V gene somatic mutations in HR2 coldspot antibodies over time. Colored dots match the clone present at all three time points in (C), empty dots indicate antibodies from other clones. Antibody hr2.016 (13) is indicated by a triangle. Mean is shown, unpaired t-test. ** is p < 0.01 and **** is p < 0.0001.

To evaluate HR2 coldspot antibody evolution at the monoclonal level, we next analyzed memory B cell antibodies derived at the later time points from two individuals with high plasma reactivity, for which we had previously obtained memory B cell antibodies at approximately 6 months (CLM41 and CLM30) (13). Peripheral blood B cells binding to the HR2 coldspot peptide were sorted as single cells by fluorescence-activated cell sorting (FACS) and the antibody sequences derived (Fig. 1B, Fig. S1B and Table S1). Although clonally related antibodies were found at all three time points from CLM41 (bright green clone with *IGHV3-15/IGKV4-1*), the overall number of V gene somatic mutations was similar over time (Fig. 1, C and D). In contrast, somatic mutations increased significantly in antibodies from CLM30, including in the clone with hr2.016, a potent and broadly neutralizing monoclonal antibody that we previously reported (bright blue clone with *IGHV3-23/IGKV3-20*; Fig. 1, C and D) (13). Other clones from this study participant included one with *IGHV1-46/IGLV1-51* (pink in Fig. 1C), which is similar to a clone that was previously reported by others (13, 21). Thus, clonally related HR2 antibodies can persist over time and, in some cases acquire additional V gene somatic mutations.

### Comparison of hr2.016 to newly obtained HR2 coldspot antibodies

To assess HR2 coldspot antibodies derived from later time points, we recombinantly expressed at least one representative monoclonal antibody per clone at different time points. In total, 19 new HR2 coldspot antibodies (7 from 18-month samples, named hr2.1xx; 12 from 30-month samples, hr2.2xx) were produced (Table S1). Sixteen out of 19 antibodies robustly bound to the S trimer of ancestral SARS-CoV-2 (half-maximal effective concentration, EC_50_ between 5.7 and 17.1 ng/mL; Fig. 2A and Table S2) and to the HR2 coldspot peptide (EC_50_ between 7.7 and 15.9 ng/mL; Fig. 2B and Table S2), which was comparable to the previously reported hr2.016 evaluated alongside. Like hr2.016, select antibodies cross-reacted with S corresponding to variants of concern (Delta, BA.1, BA.5) and Middle East respiratory syndrome coronavirus (MERS-CoV) (Fig. 2C, top, and Fig. S2A), as well as with HR2 peptides corresponding to a number of related coronaviruses (Fig. 2C, bottom, and Fig. S2B). However, none of the new antibodies displayed a substantially improved cross-reactivity profile over hr2.016.

**Figure 2.**
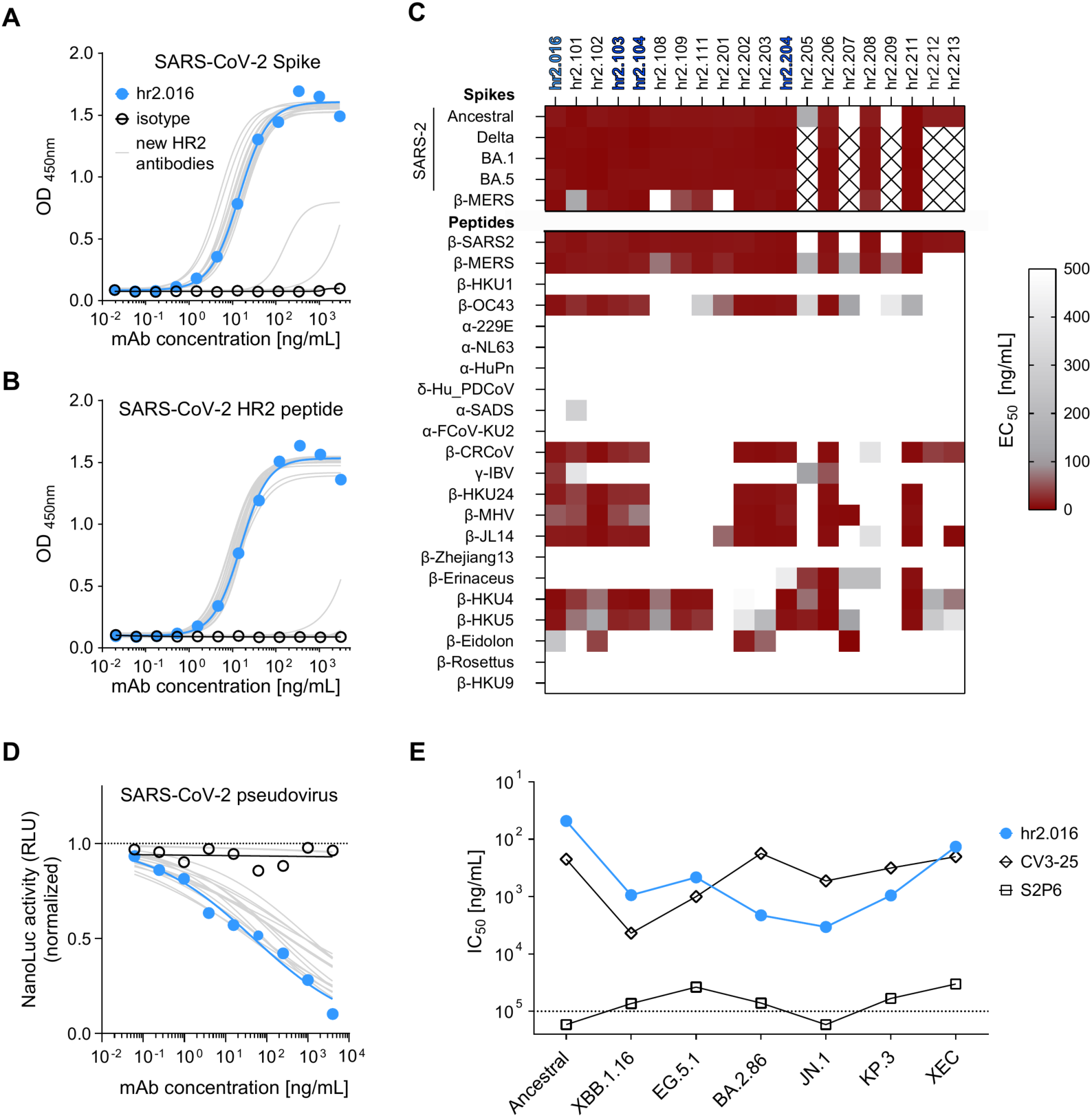
Binding and neutralizing properties of HR2 coldspot monoclonal antibodies. (**A** and **B**) ELISA measuring antibodies binding to SARS-CoV-2 ancestral S trimer **(A)** and HR2 coldspot peptide **(B)**. Mean of two independent experiments. **(C)** Heatmap displaying the summary of ELISA EC_50_ values of antibodies binding to S (top) and HR2 coldspot peptides (bottom) of the indicated coronaviruses. “X” indicates not tested. Antibodies clonally related to hr2.016 are highlighted in blue. (**D**) Normalized relative luminescence in cell lysates 48 h after infection with ancestral SARS-CoV-2 pseudovirus in the presence of increasing concentrations of HR2 coldspot antibodies. **(E)** Summary of neutralization IC_50_ values for hr2.016 in side-by-side comparison with antibodies CV3-25 (17) and S2P6 (8) against pseudoviruses corresponding to the indicated SARS-CoV-2 variants.

To evaluate the HR2 coldspot antibodies’ neutralizing capacity, we used a previously established pseudovirus system with nanoluciferase (22). Only two of the 16 new antibodies that were tested displayed half-maximal inhibitory concentrations (IC_50_) lower than 50 ng/mL against ancestral SARS-CoV-2 pseudovirus: hr2.102 and hr2.202 (39 ng/mL and 48 ng/mL, respectively), which were comparable to hr2.016 (IC_50_ of 48 ng/mL; Fig. 2D and Table S2). Similar results were obtained with pseudoviruses corresponding to virus variants (Fig. 2E, Fig. S3, A and B, and Table S2). Furthermore, hr2.016 displayed only low polyreactivity and exhibited a half-life *in vivo* of 189 h (Fig. S4). Thus, despite the acquisition of somatic mutations, none of the HR2 coldspot antibodies that were obtained at 18 or 30 months was substantially better than hr2.016, which displays suitable features for further clinical development.

### Structural comparison of HR2 stem helix antibodies with similar binding but distinct neutralizing activity

To define the structural basis of HR2 stem helix recognition, we determined crystal structures of a subset of HR2-specific antibodies that we previously isolated and characterized (13), encompassing three distinct germline usages: the recurrent public *IGHV1-46/IGKV3-20* lineage (hr2.023), the clonally related *IGHV3-23/IGKV3-20* antibodies hr2.016 and hr2.086, and an unrelated *IGHV3-74/IGKV1-4* antibody (hr2.017) (Table 1 and Fig. S5A). Despite their distinct germline origins, all four antibodies recognized the conserved hydrophobic face of the HR2 stem helix through remarkably similar overall binding modes that closely resemble those of previously described class-1 stem helix antibodies (18), supporting their classification within this antibody class. Comparison of buried surface area (BSA) further demonstrated that antibodies from these independent lineages converge on a common hotspot centered around S HR2 residues F1148, L1152, Y1155, and F1156, with slight difference in peripheral contacts (Fig. S5B), highlighting these residues as the principal energetic hotspot for HR2 stem helix recognition.

**Table 1.**
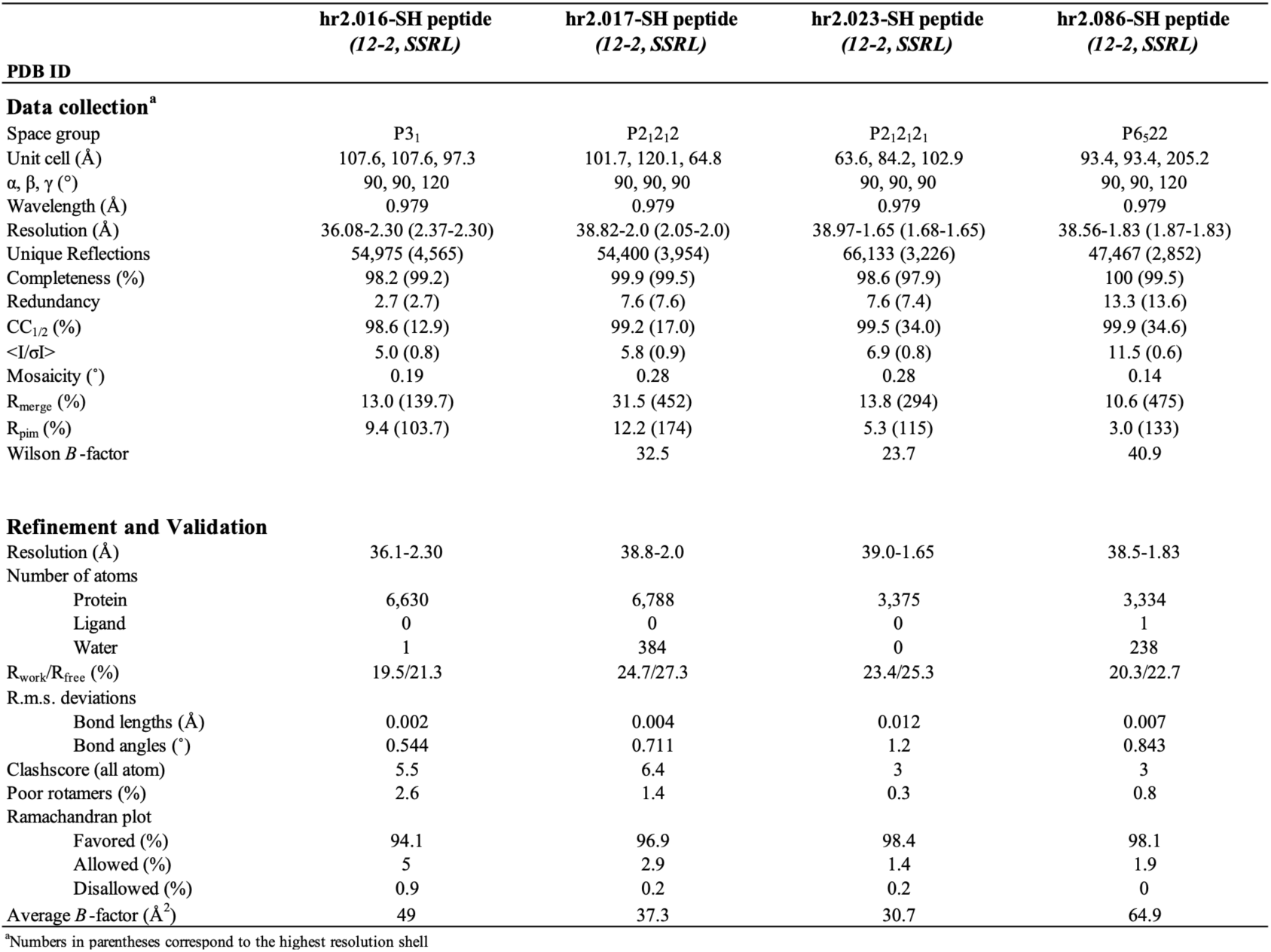
Crystallographic data collection and refinement statistics.

Among these antibodies, hr2.016 and hr2.086 were particularly intriguing because they are clonally related (*IGHV3-23*01/IGHJ4*02*; *IGKV3-20*01/IGKJ1*01*), bind similarly in ELISA, yet differ markedly in neutralization potency (Fig. 3, A and B, and in (13)). Comparison of their amino acid sequences revealed that hr2.016 and hr2.086 differed by 14 and 5 residues in the heavy and light chains, respectively, with most substitutions concentrated in the heavy-chain CDR3 and light-chain CDR1 regions (Fig. 3C).

**Figure 3.**
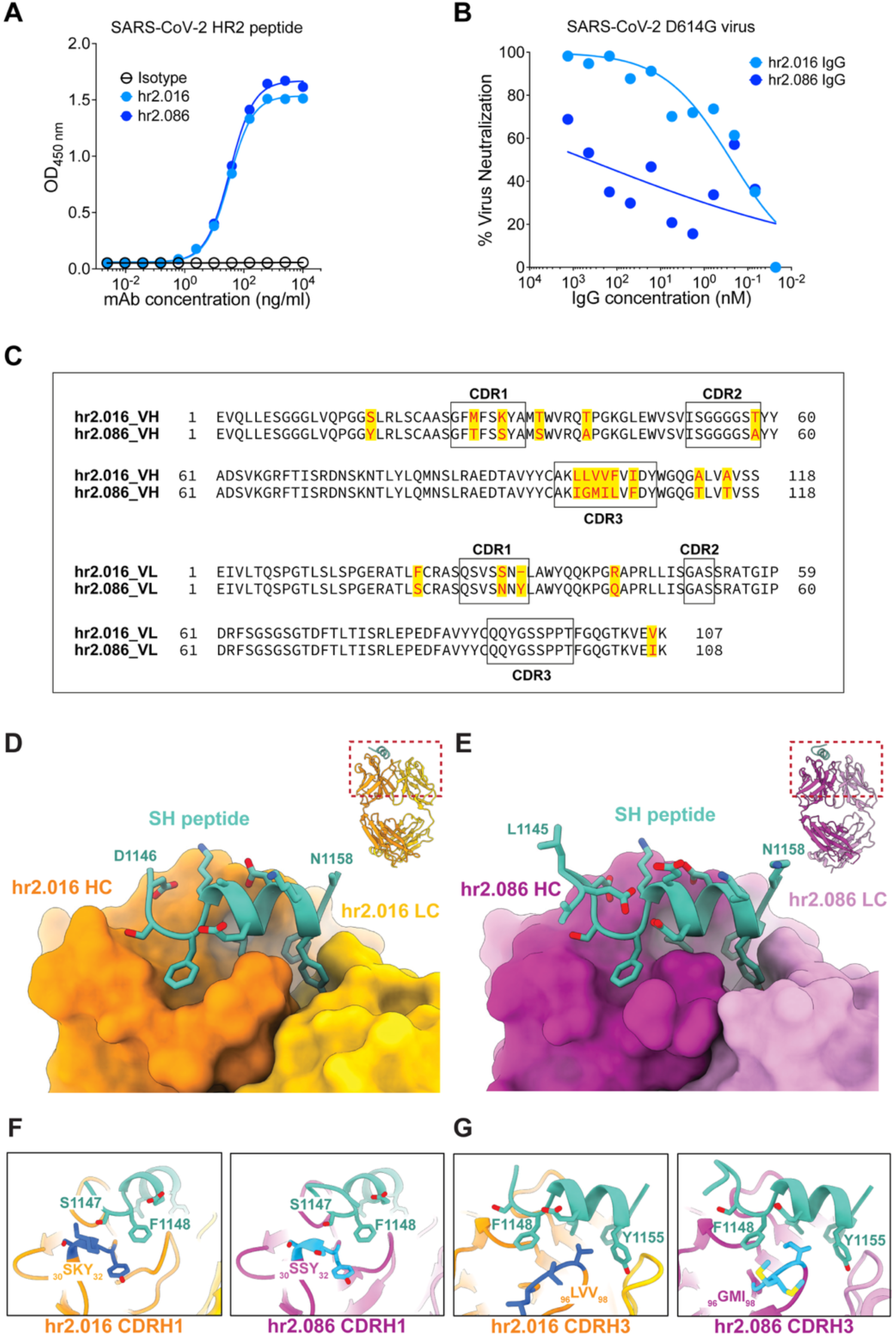
Structural comparison of clonally related HR2 coldspot antibodies with similar binding but distinct neutralizing activity. **(A)** ELISA measuring binding of hr2.016 and hr2.086 IgG to the HR2 coldspot peptide. Mean of two independent experiments**. (B)** Neutralization of SARS-CoV-2 authentic virus by hr2.016 and hr2.086. **(C)** Sequence alignment of hr2.016 and hr2.086 variable regions highlighting amino acid differences in the heavy and light chains. **(D)** Crystal structure of the hr2.016 Fab (heavy chain, orange; light chain, yellow) in complex with the HR2 coldspot peptide (teal). Inset shows the full Fab–peptide complex, with the boxed region indicating the close-up view. **(E)** Crystal structure of the hr2.086 Fab (heavy chain, magenta; light chain, pink) in complex with the HR2 coldspot peptide, shown as in **(D). (F)** Close-up view of paratope differences in CDRH1 between hr2.016 and hr2.086. **(G)** Close-up view of paratope differences in CDRH3 between hr2.016 and hr2.086.

The crystal structures of hr2.016 and hr2.086 Fabs in complex with the 20-residue HR2 coldspot peptide (^1144^ELDSFKEELDKYFKNHTSPD^1163^) were solved at 2.30 Å and 1.83 Å resolution, respectively (Fig. 3, D and E, and Table 1). In both structures, the peptide adopts an *α* -helical conformation and docks into a shallow groove formed by the heavy-chain CDRs (CDRH1/H2/H3), with CDRL3 clamping the C-terminal end of the peptide (Fig. 3, D and E). Superposition of the bound peptides (RMSD 1.47 Å over 12 Cα atoms) and similar per residue BSA values (Fig. S5B) confirm that hr2.016 and hr2.086 recognize the HR2 stem helix through essentially indistinguishable binding modes.

Consistent with their nearly identical binding geometry, PDBePISA (23) interface analysis revealed that the paratope differences between hr2.016 and hr2.086 are subtle and largely confined to CDRH1 and CDRH3, whereas the major peptide-contacting loops, including CDRH2 and CDRL3, are highly conserved (Fig. 3, F and G, and Fig. S5, C and D). Within CDRH1, the only direct paratope difference occurs at position 31, where hr2.016 contains the somatically acquired Lys31 in place of the germline-encoded Ser31 found in hr2.086 (Fig. 3F). The principal divergence resides within CDRH3, where residues 95–99 differ (LLVVF versus IGMIL), reflecting distinct V-D junction sequences; notably, Met97 of hr2.086 adopts two electron-density-supported rotamers, one of which directly contacts the peptide (Fig. 3G). Quantitatively, hr2.016 buries modestly more surface area than hr2.086 (ΔBSA ∼30 Å²), with contributions of ∼19 Å² from CDRH1 and ∼9 Å² from CDRH3 (and ∼1 Å² from CDRH2), whereas the light chain of hr2.086 buries slightly more surface (∼6 Å²). In addition, hr2.016 contains a CDRL1 deletion relative to the VK3-20 germline that although not directly contacting the peptide, subtly shifts the loop conformation and may indirectly modulate the conserved CDRL3 clamp.

Collectively, these analyses demonstrate that hr2.016 and hr2.086 engage the HR2 stem helix through nearly identical structural mechanisms despite their markedly different neutralization activities. The overall binding geometry, epitope footprint, and interaction network are highly conserved, indicating that epitope recognition alone cannot explain their divergent neutralization profiles.

### Dissociation kinetics guide SARS-CoV-2 neutralization by HR2 coldspot antibodies

Given the near-identical engagement of the epitope and the comparable binding of hr2.016 and hr2.086 IgG to the peptide and S protein in ELISA (Fig. 3A and (13)), we next examined their binding kinetics to determine whether differences in kinetic behavior might explain the disparity in neutralization potency between the two antibodies. To this end, we used surface plasmon resonance (SPR) and measured binding kinetics of either IgG or Fab, to immobilized trimeric S (Fig. 4A). IgG and Fab association rates were similar, indicating comparable efficiency of epitope binding (Fig. 4B). In contrast, the dissociation rates of the IgG were markedly different: hr2.016 exhibited a residence time of ∼4.8 hours, nearly sixfold longer than hr2.086 (∼53 minutes). Consistent with avidity effects contributing to binding, hr2.016 Fab dissociated in ∼51 minutes, closely matching hr2.086 IgG, whereas hr2.086 Fab disengaged within ∼4 minutes (Fig. 4B). The dissociation values directly paralleled pseudovirus neutralization potency, whereby only hr2.016 IgG (the format with the longest residence time), achieved robust neutralization (Fig. 4C).

**Figure 4.**
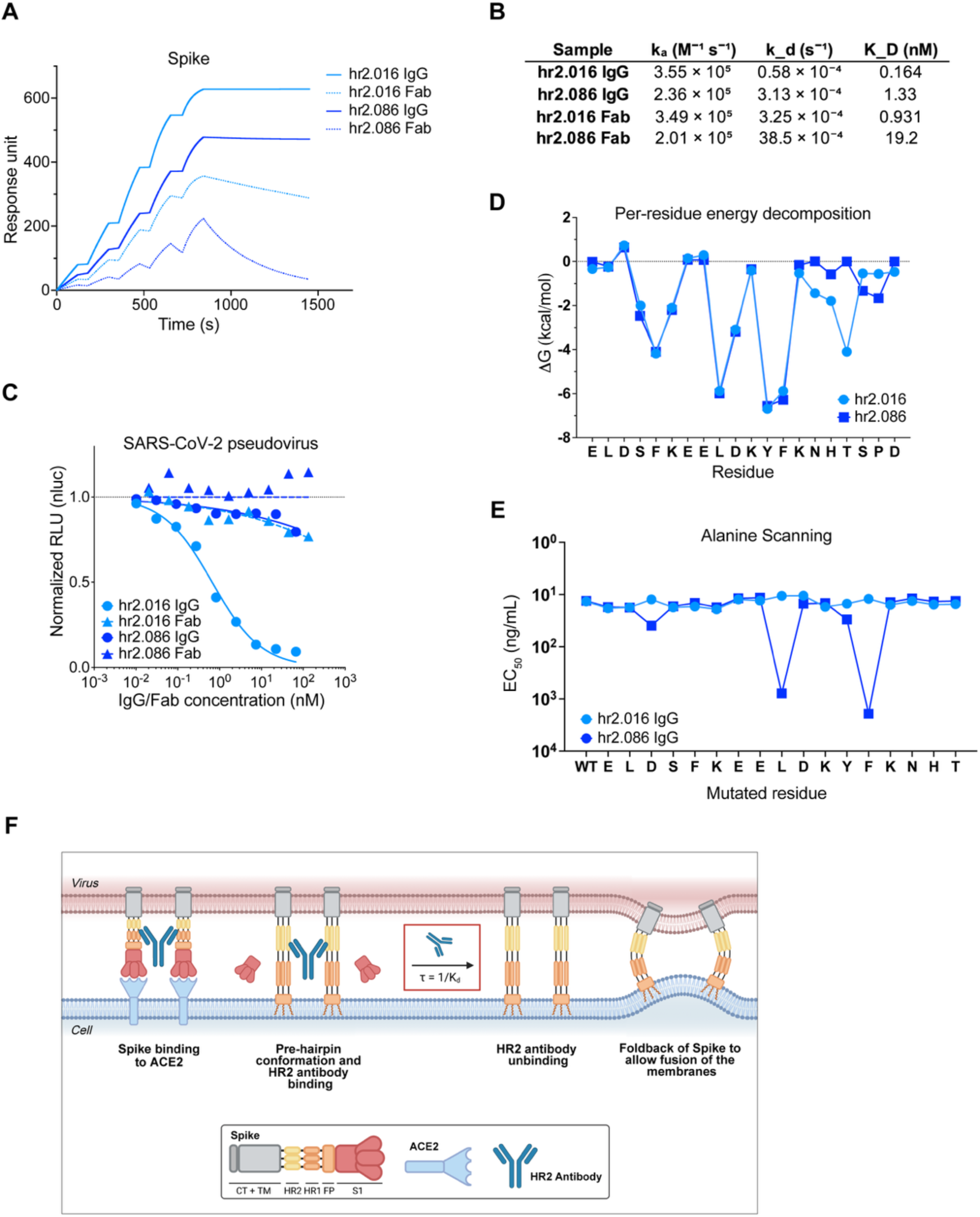
Avidity and residence time determine neutralization potency of HR2 coldspot antibodies. **(A)** Surface plasmon resonance (SPR) sensorgrams showing binding kinetics of hr2.016 and hr2.086 in IgG and Fab formats to immobilized trimeric SARS-CoV-2 S protein. Data from one representative experiment. **(B)** Association (ka) and dissociation (kd) rate constants and equilibrium dissociation constants (KD) for hr2.016 and hr2.086 IgG and Fab, determined by SPR. **(C)** Normalized relative luminescence in cell lysates 48 h after infection with ancestral SARS-CoV-2 pseudovirus in the presence of increasing concentrations of hr2.016 and hr2.086 IgG or Fab. Mean of two independent experiments. **(D)** Per-residue MM/GBSA energy decomposition (ΔG, kcal/mol) for hr2.016 and hr2.086 across the HR2 coldspot peptide, from molecular dynamics simulations. **(E)** Alanine-scanning ELISA showing EC50 values for hr2.016 and hr2.086 IgG binding to the wild-type and single-alanine-substituted HR2 coldspot peptides. **(F)** Schematic representation of antibody engagement with the HR2 stem helix on the prefusion S trimer. HR2 stem helix antibodies bind the epitope in prefusion S protein. Following ACE2 engagement, S undergoes conformational uncapping and progresses toward the fusion intermediate. Antibodies with sufficiently slow dissociation can remain bound during this transition and prevent foldback into the post-fusion conformation, thereby effectively neutralizing the virus. Created in BioRender. Guerra, C. (https://BioRender.com/v2pjj91)

To gain mechanistic insight into this kinetic divergence, we performed 1 μs molecular dynamics (MD) simulations of each Fab–peptide complex, followed by MM/GBSA calculation of binding free energies and per-residue energetic contributions, together with interaction entropy analysis. Although the two complexes converged to similar overall binding free energies, the underlying energetic balance differed markedly (Table 2). hr2.016 showed a more favorable binding enthalpy than hr2.086 (ΔH = −99.88 versus −92.78 kcal/mol), driven mainly by stronger electrostatic interactions. This enthalpic gain was counterbalanced by a larger entropic penalty, consistent with a more conformationally restricted bound state. Importantly, this entropic cost is not indicative of weaker binding, but rather of a tighter and more ordered interaction network. In kinetic terms, such an enthalpically optimized and conformationally restrained state is consistent with the prolonged residence time of hr2.016, whereas the smaller entropic penalty of hr2.086 supports a more flexible binding mode that may more readily progress toward dissociation. Per-residue energy decomposition further provided insight into the structural origin of the kinetic differences between the two antibodies (Fig. 4D). Up to approximately Phe1156, the two antibodies engaged the epitope similarly, but residues Asn1158-Thr1160 contributed additional stabilizing interactions in hr2.016 that were absent or substantially weaker in hr2.086. In particular, Thr1160 contributed −4.1 kcal/mol in hr2.016, whereas its contribution in hr2.086 was negligible, while Asn1158 and His1159 together contributed −3.2 kcal/mol. These data indicate that hr2.016 more effectively locks the peptide C-terminus into a structured interaction network, whereas the weaker stabilization by hr2.086 leaves this region more flexible and solvent-exposed. Such residual mobility likely facilitates early detachment by allowing solvent infiltration and progressive disruption of local contacts. To experimentally probe these interactions, we performed alanine scanning of the HR2 coldspot peptide and measured binding of hr2.016 and hr2.086 by ELISA (Fig. 4E). Strikingly, hr2.016 binding was largely insensitive to single alanine substitutions across the entire peptide, consistent with a distributed interaction network in which no single residue is strictly essential. In contrast, hr2.086 showed pronounced sensitivity to substitutions at select positions that correspond to peptide residue interaction hotspots. These data are consistent with the broader and more redundant engagement of the epitope by hr2.016 suggested by MD simulations, and further support the notion that a resilient, distributed interaction network, rather than reliance on discrete hotspot contacts, underlies hr2.016’s prolonged residence time and superior neutralization potency.

**Table 2.** MM/GBSA energy summary of hr2.016 and hr2.086.

**MM/GBSA energy summary – hr2.016 vs hr2.086**
| Component | hr2.016 | | hr2.086 | | $\Delta$ (hr2.086 – hr2.016) |
| --- | --- | --- | --- | --- | --- |
|  | Avg | SD | Avg | SD |  |
| MM/GBSA components |  |  |  |  |  |
| ΔVDWAALS | -89.33 | 8.56 | -84.02 | 7.44 | +5.31 |
| ΔEEL | -432.36 | 53.87 | -367.12 | 32.62 | +65.24 |
| ΔEGB | 433.96 | 52.17 | 370.10 | 30.38 | -63.86 |
| ΔESURF | -12.14 | 1.12 | -11.73 | 0.99 | +0.41 |
| Gas phase |  |  |  |  |  |
| ΔGGAS | -521.70 | 57.11 | -451.14 | 35.18 | +70.56 |
| Solvation |  |  |  |  |  |
| ΔGSOLV | 421.82 | 51.46 | 358.37 | 29.74 | -63.45 |
| Binding energy |  |  |  |  |  |
| ΔTOTAL (MM/GBSA) | -99.88 | 10.33 | -92.78 | 8.63 | +7.10 |
| Entropy (IE) |  |  |  |  |  |
| $-T\Delta$S | 120.04 | 2.53 | 107.06 | 3.24 | -12.98 |
| ΔG binding (MM/GBSA+IE) | 20.16 | — | 14.28 | — | -5.88 |

Energy values are reported in kcal/mol. $\Delta$ values are calculated as hr2.086 – hr2.016.

Together, these results indicate that the difference in neutralization between hr2.016 and hr2.086 is driven by dissociation kinetics (Fig. 4F) and reflects stronger C-terminal stabilization within a more enthalpically favorable and conformationally restrained bound state in hr2.016. Its kinetic stability, distributed interaction network, and tolerance to single substitutions make hr2.016 a remarkably resilient antibody and an especially attractive candidate for targeting the conserved viral fusion machinery of SARS-CoV-2.

## Discussion

Conserved elements of viral fusion machinery represent attractive targets for broadly reactive antibodies, yet how human responses to these epitopes evolve and what biophysical constraints govern neutralization remain poorly defined. The SARS-CoV-2 HR2 stem helix is one such site, targeted by several broadly neutralizing human antibodies (8, 11, 12, 16–18). We previously used a coldspot-guided discovery approach to identify a panel of naturally elicited antibodies against this region (13), among which hr2.016 emerged as the most promising candidate for further development. Here, we examined the long-term evolution of the HR2 coldspot response and the mechanism underlying neutralizing activity at this site. Despite clonal persistence and continued somatic hypermutation over 30 months, no HR2-specific antibody surpassed hr2.016 in combined breadth and potency, even though hr2.016 itself emerged shortly after primary infection. Structural analyses of antibodies from three distinct germline lineages revealed convergence on a common mode of stem helix recognition. Importantly, this structural convergence did not predict antiviral activity: comparison of hr2.016 with its non-neutralizing clonal relative hr2.086 identified dissociation kinetics and avidity as the decisive functional discriminators.

The limited functional improvement observed over longitudinal evolution indicates that continued somatic hypermutation does not, by itself, yield progressively greater activity at this site. This may reflect intrinsic constraints imposed by the epitope. Structural analysis of the four HR2 antibodies revealed convergence on a common hydrophobic surface, with F1148, L1152, Y1155, and F1156 forming a recognition core characteristic of class-1 stem helix antibodies (16–18).

Adopting this shared recognition mode is nonetheless not sufficient for neutralization. hr2.016 and hr2.086 derive from the same *IGHV3-23/IGKV3-20* clonotype and engage the epitope with nearly indistinguishable geometries, with differences confined to CDRH1 and CDRH3 while the major CDRH2- and CDRL3-mediated contacts are retained in both. Mutational and molecular dynamics analyses show that hr2.016 engages the peptide through a more distributed and redundant interaction network and tolerates substantially greater variation across the HR2 epitope, whereas hr2.086 depends on a narrower set of hotspot contacts. Consistently, hr2.016 Fab dissociates substantially more slowly than hr2.086 Fab, and robust neutralization is observed only for hr2.016 in the bivalent IgG format, in which dissociation is further slowed by avidity. These observations support a kinetic-gating model in which sufficiently persistent association with S allows the antibody to interfere with the conformational transitions that drive membrane fusion, highlighting residence time rather than the extent of binding as the determinant of neutralization. Similar kinetic requirements have been described for antibodies targeting the influenza hemagglutinin stem (24, 25), suggesting that this requirement may extend more broadly to antibodies directed against transiently accessible elements of viral fusion machinery.

These findings have implications for antibody discovery and immunogen design. For antibody discovery, dissociation rate and residence time may warrant priority in screening cascades when candidates share an epitope and binding geometry. For immunogen design, strategies that stabilize or otherwise favor exposure of the HR2 stem helix – for example epitope-scaffolded or nanoparticle immunogens analogous to those developed for the influenza hemagglutinin stem (26, 27) – could help overcome the natural subdominance of this site and elicit hr2.016-like responses more reliably than native S exposure alone (13).

Several limitations should be taken into account. Longitudinal monoclonal antibody analysis was restricted to two individuals with strong HR2 coldspot responses, and therefore cannot establish whether the evolutionary pattern observed here is broadly representative or reflects an absolute maturation ceiling at this epitope. In addition, although the isolated HR2 peptide enabled detailed structural characterization of antibody recognition, it may not fully reflect the steric constraints and conformational dynamics experienced by this region within membrane-embedded S.

Overall, prolonged evolution of the HR2 coldspot response did not translate into improved antibody function, and recognition of this conserved epitope alone is insufficient for neutralization. hr2.016 combines broad recognition of this structurally constrained site with the kinetic stability required to convert binding into antiviral activity. Together with its tolerance of epitope variation, good in vivo half-life and low polyreactivity, these features distinguish hr2.016 within the HR2 response and support its further development, while identifying kinetic stability as a key consideration for antibodies targeting conserved viral fusion machinery.

## Materials and Methods

### Study participants

The longitudinal COVID-19 cohort was previously described (13) and includes 71 individuals who were diagnosed with first COVID-19 infection at the Clinica Luganese Moncucco (CLM, Switzerland) between March and November of 2020. Samples for the 6 months timepoint were collected 83-269 days after onset of symptoms, later timepoints were collected at approximately 12 months intervals. The study was performed in compliance with all relevant ethical regulations under study protocols approved by the Ethical Committee of the Canton Ticino (ECCT): CE-3428 and CE-3960.

### *In vivo* mAb half-life studies

Animal experiments to determine the in vivo half-life of hr2.016 in FcγR/FcRn humanized mice were performed at The Rockefeller University and were approved by the Rockefeller University Institutional Animal Care and Use Committee in compliance with federal laws and institutional guidelines. Mice were maintained at the Comparative Bioscience Center at the Rockefeller University at a controlled ambient temperature environment with 12 h dark/light cycle. FcγR/FcRn humanized mice were generated and characterized previously (28). hr2.016 (50 μg) was administered i.v. to adult FcγR/FcR humanized mice (male and female; 7-12 week old) and blood was collected at the indicated timepoints for analysis of serum mAb levels.

Human-specific IgG ELISA was performed as previously described (29). Briefly, serum samples were serially diluted, and human hr2.016 was captured using biotinylated goat anti-human IgG, immobilized on neutravidin-coated plates and detected with HRP-conjugated anti-human IgG. After TMB development, absorbance was measured at 450 nm, and hr2.016 concentrations and serum half-life were calculated as described (30).

### Blood sample processing and storage

Peripheral blood mononuclear cells (PBMCs) were obtained by Histopaque density centrifugation, aliquoted and stored in liquid nitrogen in the presence of 10% fetal bovine serum (FBS) and dimethyl sulfoxide (DMSO). Anticoagulated plasma was aliquoted and stored at −20°C or less. Before use, aliquots of plasma were heat-inactivated (56°C for 1 hour) and then stored at 4°C.

### Peptides and recombinant proteins for biochemical studies

#### Synthetic peptides

All peptides with HR2 coldspot sequences used in this study were previously described (13), with the exception of the Alanine mutants (Fig. 4E). In all cases, peptides were biotinylated (Biotin-Ahx) at the N-terminus and amidated at the C-terminus and were obtained from GenScript (Hong Kong) with at least 75% purity.

#### Recombinant proteins

The coronaviruses S proteins were produced and purified as described (13, 31). Briefly, a codon-optimized gene encoding residues 1-1208 of SARS-CoV-2 S ectodomain (GenBank: MN908947) was synthesized and cloned into the expression vector pcDNA3.1(+) by Genscript; the sequence contains proline substitutions at residues 986 and 987 (S-2P), a ‘GSAS’ substitution at the furin cleavage site (residues 682–685), a C-terminal T4 fibritin trimerization motif, and a C-terminal octa-histidine tag for purification. SARS-CoV-2 S ectodomains corresponding to the SARS-CoV-2 VOC were similarly modified and based on: Delta, GenBank: QWK65230.1; Omicron BA.1 GenBank: UFO69279.1; Omicron BA.5 GenBank: UPP14409.1 + G3V. MERS S ectodomain was based on PDB: 6NB3_A for MERS. All proteins were produced by transient transfection of Expi293F cells (ThermoFisher) using polyethylenimine (PEI), purified from the cells supernatants with HiTrap Chelating HP (Cytiva) affinity columns and analyzed to ensure functionality, stability, lack of aggregation and batch-to-batch reproducibility as previously described (13).

### Enzyme-linked immunosorbent assays (ELISA)

#### Peptide ELISA

384-well plates (ThermoFisher, 464718) were coated with 10 μL per well of a 2 μg/mL Neutravidin (Life Technologies, 31000) solution in PBS, overnight at room temperature. Plates were washed 3 times with washing buffer (PBS + 0.05% Tween-20, Sigma-Aldrich) and incubated with 10 μL per well of a 50 nM biotinylated peptide solution in PBS for 1 h at room temperature. After washing 3 times with washing buffer, plates were incubated with 50 μL per well of blocking buffer (PBS + 1% BSA + EDTA 1 mM + 0.05% Tween-20) for 2 h at room temperature. Plates were then washed 3 times with washing buffer, and serial dilutions of plasma or monoclonal antibodies were added in PBS + 0.05% Tween-20 and incubated for 1 h at room temperature. To screen for the presence of anti-HR2 coldspot peptide IgGs, plasma samples were assayed at 1:50 starting dilution, followed by 3 three-fold serial dilutions. Monoclonal antibodies were tested starting at 3 μg/mL, followed by three-fold serial dilutions. Plates were subsequently washed 3 times with washing buffer and incubated with anti-human IgG secondary antibody conjugated to horseradish peroxidase (HRP) (GE Healthcare, NA933) at a 1:5000 dilution in PBS + 0.05% Tween-20. Finally, after washing 3 times with washing buffer, plates were developed by the addition of 10 μl per well of the HRP substrate TMB (ThermoFisher, 34021) for 10 min. The developing reaction was stopped with 10 μl per well of a 1M H2SO4 solution, and absorbance was measured at 450 nm with an ELISA microplate reader (BioTek) with Gen5 software. The background reactivity of plasma samples on neutravidin was subtracted from the measured values. The Area Under the Curve (AUC) was calculated from two independent experiments and plotted with GraphPad Prism (software v.11). The half-maximal effective concentration of monoclonal antibodies (EC_50_) was determined using four-parameter nonlinear regression curve fit (GraphPad Prism, software v.11).

#### Protein ELISA

Experiments were performed with 384-well plates coated with 10 μL per well of a 5 μg/mL protein solution in PBS overnight at room temperature and subsequently blocked and treated as described above. Monoclonal antibodies were tested starting at 3 μg/mL, followed by threefold serial dilutions.

#### ELISA for antibody polyreactivity

Antibody polyreactivity was assessed against ssDNA, dsDNA, LPS, KLH, and insulin, following a previously described protocol (29). Briefly, antigen-coated plates were blocked and incubated with monoclonal antibodies for 1 h, followed by detection with the same secondary antibody and development conditions described above. The polyreactive antibody ED38 was used as a positive control (32), while the ZIKV-specific antibody Z021 served as a negative control (33).

### Single-cell sorting by flow cytometry

HR2 coldspot peptide-specific B cells were derived from PBMCs of COVID-19 convalescent individuals as previously detailed (13). Briefly, B lymphocytes were enriched using the pan-B-cell isolation kit according to manufacturer’s instructions (Miltenyi Biotec, 130-101-638) and stained in FACS buffer (PBS + 2% FCS + 1 mM EDTA) with the following antibodies/reagents (all at 1:200 dilution) for 30 min on ice: anti-CD20-PE-Cy7 (BD Biosciences, 335828), anti-CD14-APC-eFluor 780 (Thermo Fischer Scientific, 47-0149-42), anti-CD16-APC-eFluor 780 (Thermo Fischer Scientific, 47-0168-41), anti-CD3-APC-eFluor 780 (Thermo Fischer Scientific, 47-0037-41), anti-CD8-APC-eFluor 780 (Invitrogen, 47-0086-42), Zombie NIR (BioLegend, 423105), as well as fluorophore-labeled ovalbumin (Ova) and peptides. Live single Zombie-NIR−CD14−CD16−CD3−CD8−CD20+Ova−peptide-PE+peptide-AF647+ B cells were single-cell sorted into 96-well plates containing 4 μl of lysis buffer (0.5× PBS, 10 mM DTT, 3,000 units/ml RNasin Ribonuclease Inhibitors, Promega, N2615) per well using a FACS Aria III, and the analysis was performed with FlowJo software. The gating strategy is shown in Fig. S1B.

### Antibody gene sequencing, cloning and expression

Antibody gene sequences were identified as described previously (13). Briefly, single cell RNA was reverse-transcribed (SuperScript III Reverse Transcriptase, Invitrogen, 18080-044) and the cDNA stored at −20°C or used immediately for subsequent nested PCR amplification of the expressed variable IGH, IGL and IGK genes and Sanger sequencing. Cloning of antibody genes into expression vectors was by Sequence- and Ligation-Independent Cloning (SLIC). Recombinant monoclonal antibodies and Fabs were produced and purified as previously described (13). hr2.016 anti-SARS-CoV-2 antibody and isotype control antibody Z021 were previously published (13, 33). All recombinant antibodies were produced in Expi293 cells, purified from clarified supernatant using protein G (Protein G Sepharose™ 4 Fast Flow, Cytiva) or Ni-IMAC resin (Pierce™ High-Capacity EDTA-Compatible Ni-IMAC Resin, Thermo Fisher Scientific) and tested to ensure functionality, stability and batch-to-batch reproducibility before use in experiments.

### SARS-CoV-2 pseudotyped reporter viruses

The generation of plasmids to express a C-terminally truncated SARS-CoV-2 S protein (pSARS-CoV2-S_trunc_), the HIV-1 structural/regulatory proteins (pHIV_NL_GagPol) and the NanoLuc reporter construct (pCCNanoLuc2AEGFP) were previously described (22). These plasmids, including the pSARS-CoV2-S_trunc_ for Delta variant, were kindly gifted by Dr. Paul Bieniasz and Dr. Theodora Hatziioannou (The Rockefeller University, New York). Plasmids expressing Omicron XBB.1.16, EG.5.1, BA.2.86, JN.1, KP.3 and XEC SARS-COV-2 S_trunc_ variants were generated in house by site-directed mutagenesis (QuikChange Multi Site-Directed Mutagenesis Kit, Agilent) starting from Omicron BA.1 SARS-CoV-2 S_trunc_ in pcDNA3.1(+) (Genscript). The sequences corresponding to SARS-CoV-2 variants were based on: Omicron XBB.1.16, (GenBank UZG29433.1 + K474R); Omicron EG.5.1 (GenBank WGM84363.1); Omicron BA.2.86 (GenBank WRK62249.1 + N17M + N17_L18insPLFN + I670V); Omicron JN.1 (WRK13149.1); Omicron KP.3 (WRK13149.1 + F456L + Q493E + V1104L) and Omicron XEC (WRK13149.1 + T22N + F59S + F456L + Q493E + V1104L). In all plasmids, the DNA encoding for the intracellular domain was similarly truncated. The generation of pseudotyped virus stocks was done as previously described(13, 22). Briefly, 293T cells were transfected with pHIVNLGagPol, pCCNanoLuc2AEGFP and pSARS-CoV2-Strunc plasmids using PEI-MAX (Polysciences). At 24 h after transfection, supernatants containing non-replicating virions were harvested, filtered and stored at −80°C. Infectivity was determined by titration on 293T_ACE2_ cells.

### Pseudotyped virus neutralization assay

The assay was performed as previously described (22). Briefly, three- or fourfold serially diluted monoclonal antibodies were incubated with the SARS-CoV-2 pseudotyped virus for 1 hour at 37°C degrees. The mixture was subsequently incubated with 293T_ACE2_ cells for 48 hours, after which cells were washed once with PBS and lysed with Luciferase Cell Culture Lysis 5x reagent (Promega). NanoLuc Luciferase activity of lysates was then measured using the Nano-Glo Luciferase Assay System (Promega) with GloMax Discover System reader (Promega). Relative luminescence units were normalized to those derived from cells infected with SARS-CoV-2 pseudotyped virus in the absence of monoclonal antibodies. The half-maximal inhibitory concentration of monoclonal antibodies (IC_50_) was determined using four-parameter nonlinear regression curve fit (GraphPad Prism, software v.11).

### Real Virus Experiments

#### Virus Stocks and Titration (Focus Forming Assay)

The SARS-CoV-2 D614G strain originated from hCoV-19/France/GE1973/2020 and was kindly provided by the National Reference Center for Respiratory Viruses hosted by the Institut Pasteur and headed by Prof. Sylvie van der Werf. The virus was amplified in Vero E6 cells. Virus titration was performed by focus forming assay as previously described (34). Briefly, serial 10-fold dilutions of the virus, prepared in DMEM GlutaMAX containing 1% FBS, were used to infect confluent Vero E6 cells monolayers for 2 h at 37°C, 5% CO₂. The inoculum was removed, and cells were overlaid with semi-liquid medium consisting of MEM 1X, 1.5% CMC, 10% FBS. Plates were incubated for 48 h at 37°C, 5% CO₂. Cells were fixed with 4% formaldehyde, and viral foci were detected using rabbit anti-SARS-CoV-2 N primary antibody, followed by HRP-conjugated secondary antibody and DAB staining. Foci were counted with the Immunospot (CTL) and viral titers were expressed as focus-forming units per milliliter (FFU/ml).

#### Real Virus Neutralization Experiments

For neutralization assay, Vero E6 cells were seeded a day prior to infection at 2 × 10^4^ cells per well in DMEM GlutaMAX supplemented with 10% FBS in a 96-well plate. Virus stocks were diluted to a final concentration of 2000 FFU/ml, corresponding to 200 FFU/well (final concentration). Virus was incubated for 1 hour at 37°C in a 1:1 ratio with IgG hr2.016 or hr2.086 separately, starting from a 1.33 µM antibody concentration followed by 1:3 serial dilutions. Virus incubated with medium in the absence of antibodies was included as a control and for later normalization. The cell supernatant was removed and replaced with 100 µl of virus-antibody solution and incubated for 2 h at 37°C, 5% CO₂. The inoculum was removed and cells were overlaid with CMC containing semi-liquid medium. 48 h post infection, cells were fixed and foci were revealed as described for the focus forming assay.

#### Surface plasmon resonance (SPR)

Binding kinetics of hr2.016 and hr2.086 (both IgG and Fab) were measured on a Biacore 8K instrument (Cytiva) at 25°C using CM5 sensor chips (Cytiva). Prefusion trimeric SARS-CoV-2 Spike ectodomain was immobilized by amine coupling to a level of 6000 RU. Antibodies were injected in running buffer (10 mM HEPES pH 7.4, 150 mM NaCl, 3 mM EDTA, 0.05% Tween20) at a flow rate of 30 μL/min using a single-cycle format with a 5-point concentration series (50, 25, 12.5, 6.25, 3.12 nM). Association and dissociation were monitored for 120s (5 cycles) and 7200 s, respectively. Sensorgrams were double-referenced (reference flow cell and buffer blank) and globally fit using a 1:1 model in Biacore Evaluation Software 5.0.18 to obtain k_on_, k_off_, and KD. Residence time was calculated as 1/k_off_.

#### Molecular dynamics (MD) simulations

Starting coordinates for hr2.016-HR2 peptide and hr2.086-HR2 peptide complexes were taken from the corresponding crystal structures. Missing side chains were modeled using ALMOST (35). Each complex was solvated within a cubic periodic box of TIP3P water molecules (36), ensuring a minimum buffer distance of 10 Å between the solute atoms and the boundary of the simulation cell. To ensure the electrostatic stability of the system, charge neutrality was achieved by the addition of counterions. The ff14SB (37) force field was used to describe the protein while ion parameters were taken from Joung et al. (38) All MD simulations were performed under isothermal-isobaric conditions. The system temperature was maintained at 298.5 K using a Langevin thermostat (39), while the pressure was regulated at 1 atm via a Monte Carlo barostat (40). Long-range electrostatic interactions were treated using the Particle Mesh Ewald (PME) method.

Production runs were extended to a cumulative simulation time of 1 μs. For subsequent trajectory analysis, a total of 1,250 snapshots were extracted at uniform intervals throughout the simulation. All MD trajectories were generated using the GPU-optimized implementation of the AMBER24 package, leveraging CUDA acceleration to achieve the 1 μs production timescale. To evaluate the thermodynamic stability of the complexes, the protein-peptide binding free energies were estimated using the MM/GBSA (Molecular Mechanics Generalized Born Surface Area) method as implemented in gmx_MMPBSA (41).

In particular, the total binding free energy was estimated as ΔG = *Δ* H-T *Δ* S. *Δ* H = ΔE_VdW_+ ΔE_ele_ + ΔG_polar_ + ΔG_nonpolar_, where ΔG_polar_ was computed with a generalized Born model igb=8, salt concentration 0.150M. Reported values represent mean ± s.d. across frames. To account for entropic changes upon binding, the conformational entropy loss (TΔS) was calculated using the last 325 frames and applying the Interaction Entropy (IE) method (42).

#### X-ray crystallography

X-ray crystallography experiments were performed as previously described (13) with modifications described below. Purified Fabs were mixed with a 20-residue HR2 stem helix peptide (^1144^ELDSFKEELDKYFKNHTSPD^1163^) at a 1:2 molar ratio (Fab:peptide) and incubated overnight at room temperature. Fab–peptide complexes were concentrated using Amicon centrifugal filters with a 30-kDa molecular weight cutoff (MilliporeSigma) to 10-15 mg/mL. Crystallization trials were performed by sitting-drop vapor diffusion by mixing equal volumes of Fab–peptide complex and reservoir solution using a Mosquito LCP robot (SPT Labtech) and commercially available 96-well crystallization screens (Hampton Research). Crystals were grown at 16°C. Crystals used for structure determination were obtained under the following conditions: hr2.016, 1.5 M ammonium phosphate dibasic, 0.1 M Tris pH 8.5; hr2.086, 2 M ammonium sulfate, 0.1 M sodium acetate pH 4.5; hr2.017, 0.2 M magnesium chloride hexahydrate, 0.1 M HEPES pH 7.5, 25% polyethylene glycol 3,350; and hr2.023, 0.1 M sodium citrate tribasic dihydrate pH 5.5, 18% polyethylene glycol 3,350. Crystals were cryoprotected in complex-specific cryoprotection conditions and cryocooled in liquid nitrogen. X-ray diffraction data were collected at the Stanford Synchrotron Radiation Lightsource (SSRL) beamline 12-2 using an Eiger X 16M detector (Dectris) at a wavelength of 0.979 Å and a temperature of 100 K. Diffraction data were indexed and integrated using XDS (43) or DIALS (44) and merged and scaled using AIMLESS (45) within the CCP4 (46) suite. Structures were determined by molecular replacement in PHASER (47) using appropriate antibody variable and constant domains as search models. Coordinates were refined through iterative rounds of automated refinement in Phenix (48) and manual model building in Coot (49). Data collection and refinement statistics for all structures are provided in Table 1.

### Structural Analyses

CDR and somatic mutation assignments were produced by IMGT V-QUEST (50). Graphics describing structures were made in ChimeraX (51). Buried surface areas were calculated using the online PDBePISA server (23). Contacting residues are defined as those with less than 4 Å distance between atoms of different chains. Hydrogen bond assignments were made using a 3.5 Å cutoff and A-D-H angle greater than 90°. RMSD calculations were done in PyMOL (Schrödinger). Antibody residues are numbered according to the Kabat convention.

## Acknowledgments

We thank the study participants and their families, as well as the medical personnel at the Clinica Luganese Moncucco. We further thank Theodora Hatziioannou and Paul Bieniasz (Rockefeller University) for sharing plasmids and protocols for SARS-CoV-2 pseudovirus. At the IRB, we are grateful to V. Cecchinato and M. Uguccioni (Human Subject Research). The work was supported in part by the Swiss Vaccine Research Institute (SVRI), George Mason University Fast Grant, and NIH grants U01 AI151698 (United World Antiviral Research Network, UWARN), P01 AI138938, and U19 AI111825 (to D.F.R.); European Union’s Horizon 2020 research and innovation programme under grant agreement no. 101003650: Antibody Therapy Against Coronavirus - ATAC (to L.V. and D.F.R.); and BRIDGE 40B2-0_203488 (to A.C. and D.F.R.). Further support was provided by the Swiss National Science Foundation and in part by funds from the Howard Hughes Medical Institute Emerging Pathogens Initiative (C.O.B.). Additionally, C.O.B. is supported by the Rita Allen Foundation, Pew Biomedical Scholars Program, and is Freeman Hrabowski Scholar (HHMI). This article is subject to HHMI’s Immediate Access to Research policy, which requires that this article be made publicly available as initial and revised preprints deposited on a designated preprint server under a CC BY 4.0 license. Use of the Stanford Synchrotron Radiation Lightsource, SLAC National Accelerator Laboratory, is supported by the U.S. Department of Energy, Office of Science, Office of Basic Energy Sciences under Contract No. DE-AC02-76SF00515. The SSRL Structural Molecular Biology Program is supported by the DOE Office of Biological and Environmental Research, and by the National Institutes of Health, National Institute of General Medical Sciences (P30GM133894). The contents of this publication are solely the responsibility of the authors and do not necessarily represent the official views of NIGMS or NIH. The study was also possible thanks to the IRB-Rockefeller University partnership for infectious disease research, supported in part by a grant to the IRB from the Fondazione Leonardo.

## Data and materials availability

All data supporting the findings of this study are available in the main text or the Supplementary Information. The heavy- and light-chain sequences of the newly reported HR2 coldspot antibodies (hr2.1xx and hr2.2xx series) are provided in Table S1. The atomic models and corresponding structure factors for all structures reported in this study will be available in the Protein Data Bank (PDB) upon publication. The hr2.016–HR2 stem helix, hr2.086–HR2 stem helix, hr2.017–HR2 stem helix, and hr2.023–HR2 stem helix complexes will be available under accession codes 37KZ, 37LC, 37LA, and 37LB, respectively.

Requests for materials should be addressed to the corresponding author A.C., and may require a completed material transfer agreement.

## Usage of Generative AI

The authors declare the use of generative AI in the writing process. According to the GAIDeT taxonomy (2025), the following tasks were delegated to GAI tools under full human supervision:

- Proofreading and editing

The GAI tools used were: Chat GPT 5.6, Claude Sonnet 5.

Responsibility for the final manuscript lies entirely with the authors.

GAI tools are not listed as authors and do not bear responsibility for the final outcomes.

Declaration submitted by: The authors

## Code availability

No custom code was used in this study. Data analysis was performed using previously published or commercially available software, including GraphPad Prism (v.11), Biacore Evaluation Software (v.5.0.18), the AMBER24 simulation package, and the ProDy toolkit, as described in the Methods.

---

**Fig. S1.**
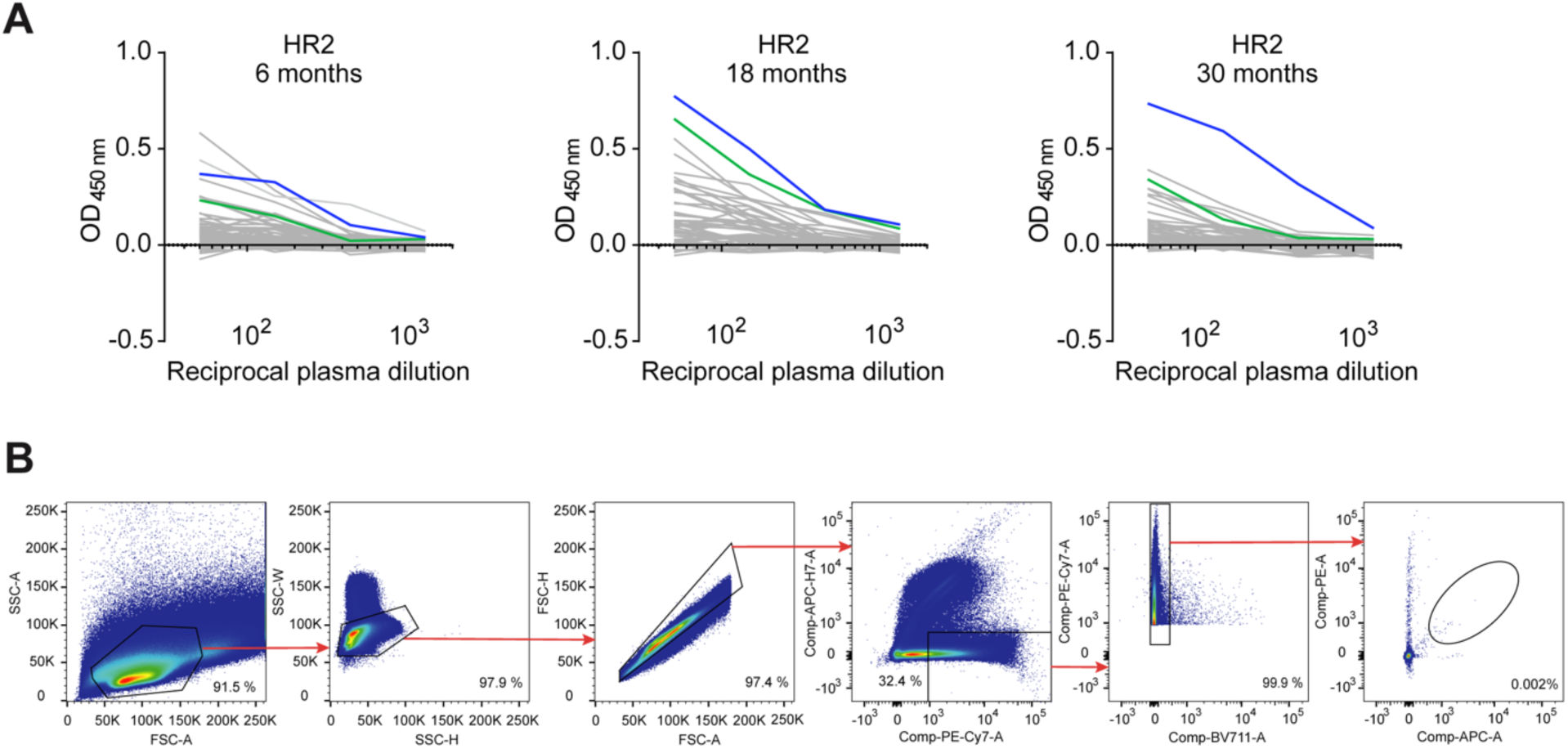
Longitudinal analysis of HR2 coldspot antibody response. (**A**) ELISAs measuring plasma IgG reactivity to the HR2 coldspot peptide at 6, 18 and 30 months after SARS-CoV-2 infection. Data are shown as mean of two independent experiments. Green (CLM41) and blue (CLM30) represent samples from individuals selected for HR2 coldspot antibody analysis. (**B**) Gating strategy used to sort HR2 coldspot peptide-specific B cells by flow cytometry.

**Fig. S2.**
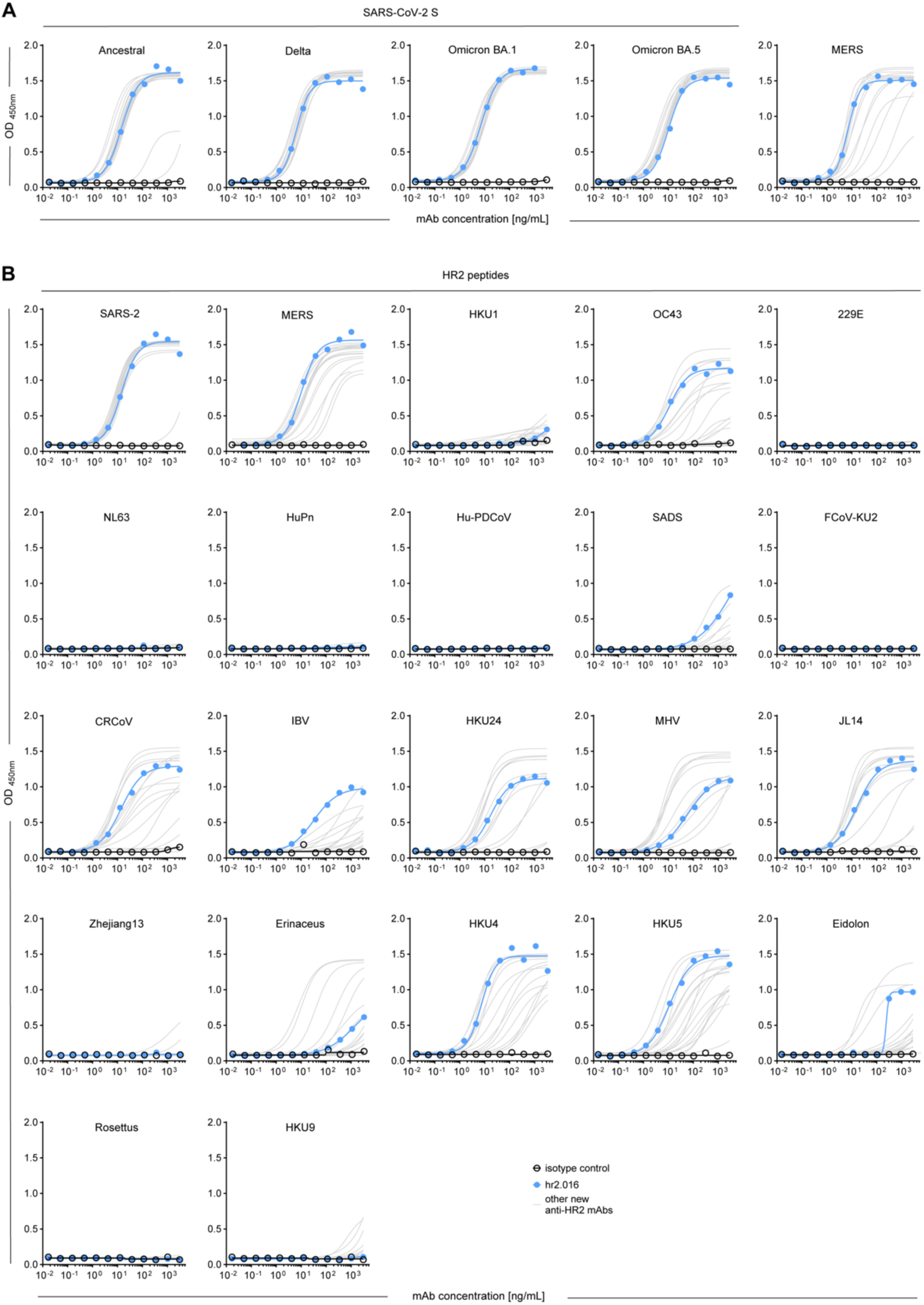
Crossreactivity of the newly isolated HR2 coldspot antibodies. (**A**) and (**B**) ELISAs measuring HR2 coldspot antibody reactivity to S proteins (A) and HR2 coldspot peptides (B) from different coronaviruses. Data are shown as the mean of two independent experiments.

**Fig. S3.**
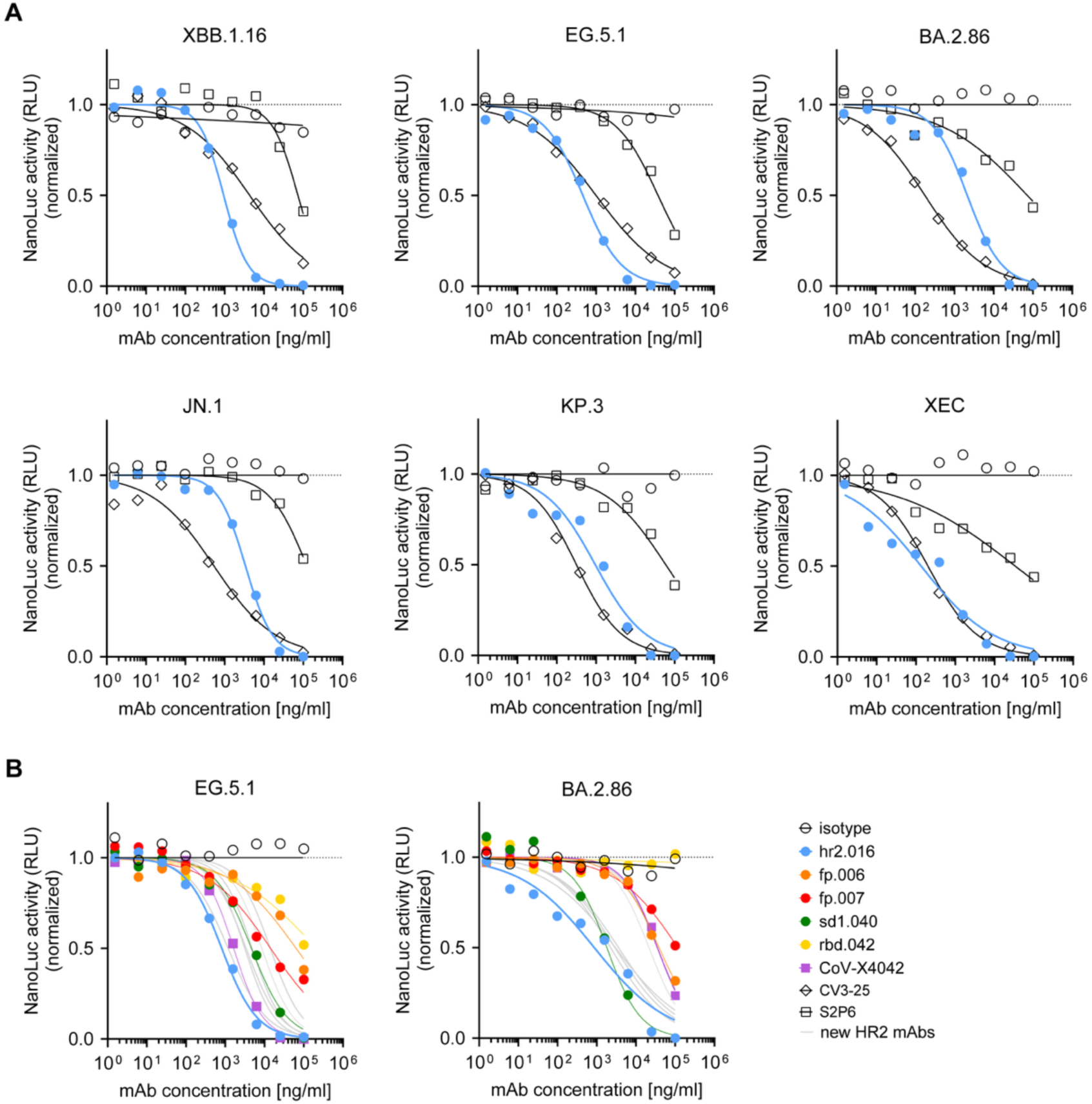
Broad neutralizing activity of hr2.016. (**A**) Graphs show normalized relative luminescence values measured in cell lysates 48 hours after infection with pseudoviruses of SARS-CoV-2 variants of concern (VOCs) in the presence of increasing concentrations of monoclonal antibodies. An isotype control and the HR2 stem helix antibodies CV3-25 and S2P6 were included for comparison. Data are shown as the mean of two independent experiments. (**B**) Same as (A) but including previously published FP, RBD and SD1 antibodies alongside the newly isolated HR2 coldspot antibodies. Data are shown as the mean of two independent experiments.

**Fig. S4.**
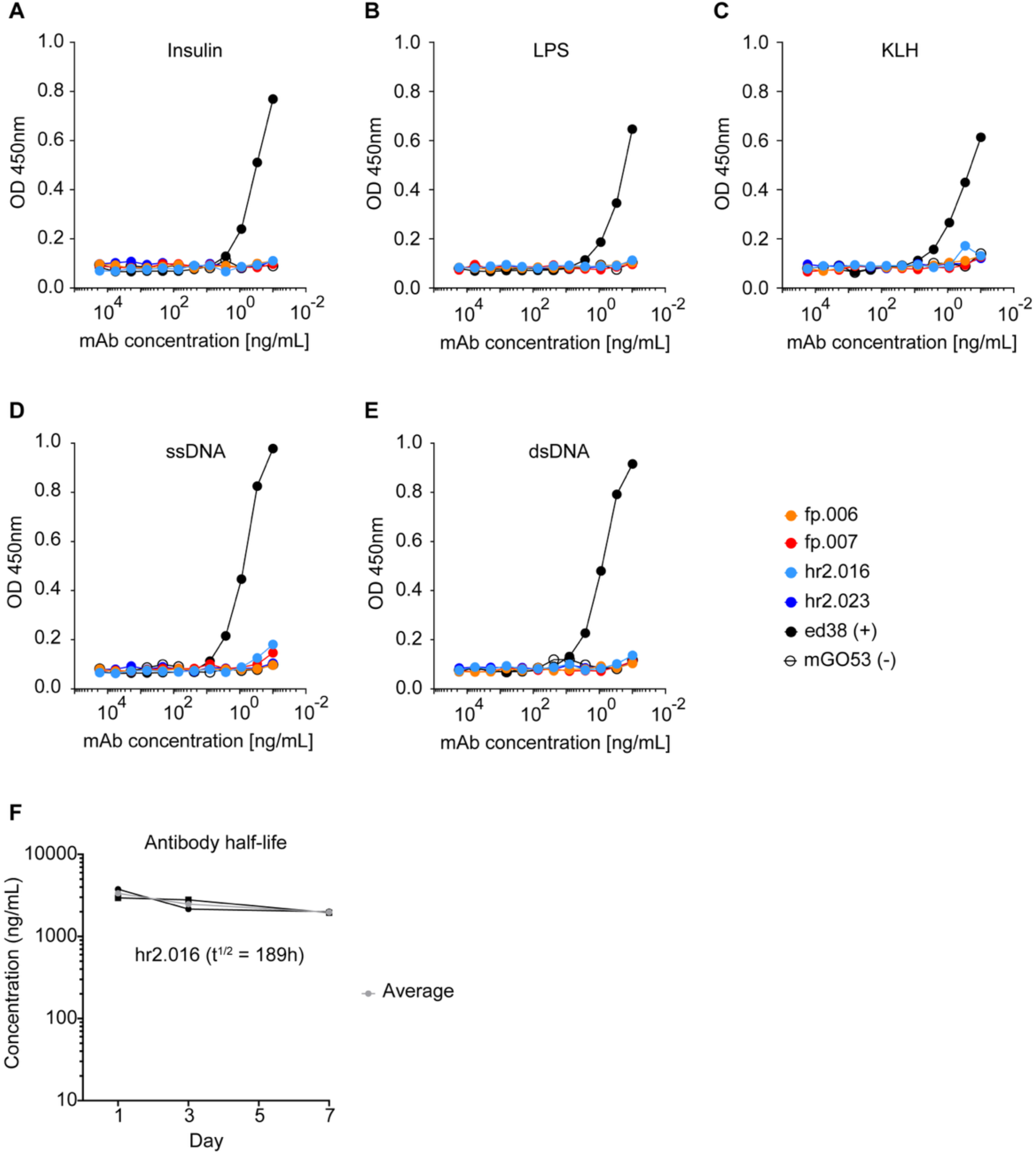
Developability properties of hr2.016. **(A**) Binding of hr2.016 to insulin, lipopolysaccharide (LPS), keyhole limpet hemocyanin (KLH), single-stranded DNA (ssDNA), and double-stranded DNA (dsDNA), evaluated by ELISA. ED38, a polyreactive monoclonal antibody, was used as a positive control. fp.006, fp.007, and hr2.023 were included for comparison. (**B**) Half-life of hr2.016 in humanized mice following intravenous administration.

**Fig. S5.**
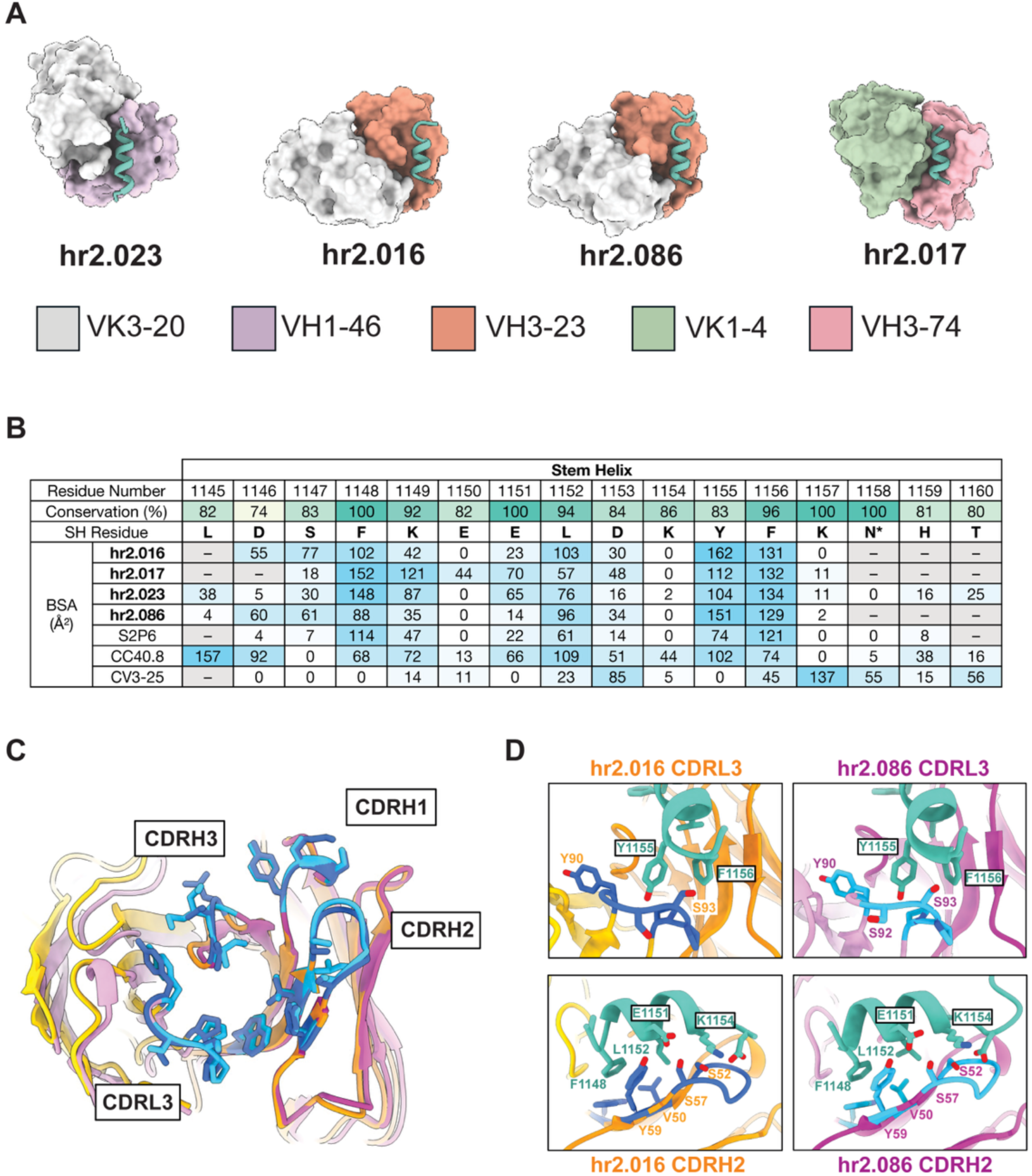
Structural convergence of antibodies targeting the HR2 stem helix. **(A)** Crystal structures of four HR2 stem helix-directed antibodies representing distinct germline solutions. Heavy- and light-chain germline gene usage is indicated. **(B)** Buried surface area (BSA) contributed by individual HR2 stem helix residues at the antibody interface for hr2.016, hr2.017, hr2.023, and hr2.086, compared with previously characterized stem helix antibodies S2P6, CC40.8, and CV3-25. Sequence conservation at each HR2 residue across betacoronaviruses is indicated above the heatmap. **(C)** Comparison of hr2.016 and hr2.086 bound to the HR2 stem helix peptide. Paratopes are shaded as blue, with selected antibody residues contributing to the interface shown as sticks. **(D)** Comparison of the CDRL3 (top panels) and CDRH2 (bottom panels) loops of hr2.016 and hr2.086, demonstrating conservation of the principal peptide-contacting residues and interactions between the two clonally related antibodies. Selected interacting residues are shown as sticks.

## Notes

### Summary of Updates

This revised version replaces the previous preprint. The main text and figures have been reformatted to improve legibility, with no changes to the data, analyses, or conclusions. The author list has also been updated to accurately reflect contributions to the work; all authors, including those added and removed, have seen and approved this revised version and the updated authorship.

